# Rare and population-specific functional variation across pig lines

**DOI:** 10.1101/2022.02.01.478603

**Authors:** Roger Ros-Freixedes, Bruno D. Valente, Ching-Yi Chen, William O. Herring, Gregor Gorjanc, John M Hickey, Martin Johnsson

## Abstract

**Background:** It is expected that functional, mainly missense and loss-of-function (LOF), and regulatory variants are responsible for phenotypic differences among breeds, genetic lines, and varieties of livestock and crop species that have undergone diverse selection histories. However, there is still limited knowledge about the existing missense and LOF variation in livestock commercial populations, in particular regarding population-specific variation and how it can affect applications such as across-breed genomic prediction.

**Methods:** We re-sequenced the whole genome of 7,848 individuals from nine commercial pig breeding lines (average sequencing coverage: 4.1x) and imputed whole-genome genotypes for 440,610 pedigree-related individuals. The called variants were categorized according to predicted functional annotation (from LOF to intergenic) and prevalence level (number of lines in which the variant segregated; from private to widespread). Variants in each category were examined in terms of distribution along the genome, minor allele frequency, Wright’s fixation index (F_ST_), individual load, and association to production traits.

**Results:** Of the 46 million called variants, 28% were private (called in only one line) and 21% were widespread (called in all nine lines). Genomic regions with low recombination rate were enriched with private variants. Low-prevalence variants (called in one or a few lines only) were enriched for lower allele frequencies, lower F_ST_, and putatively functional and regulatory roles (including loss-of-function and deleterious missense variants). Only a small subset of low-prevalence variants was found at intermediate allele frequencies and had large estimated effects on production traits. Individuals on average carried less private deleterious missense alleles than expected compared to other predicted consequence types. A small subset of low-prevalence variants with intermediate allele frequencies and higher F_ST_ were detected as significantly associated to the production traits and explained small fractions of phenotypic variance (up to 3.2%). These associations were tagged by other more widespread variants, including intergenic variants.

**Conclusions:** Most low-prevalence variants are kept at very low allele frequency and only a small subset contributed detectable fractions of phenotypic variance. Not accounting for low-prevalence variants is therefore unlikely to hinder across-breed analyses, in particular for genomic prediction of breeding values using reference populations of a different genetic background.

## Introduction

Genetic variation is the basis of selective breeding in livestock and crop species. From a molecular point of view, genetic variants that result in either altered protein structures or altered gene expressions are believed to be responsible for much of the existing genetic variation in complex traits [1–4]. Missense variants change one amino acid of the encoded protein. Loss-of-function variants (LOF) are predicted to disrupt protein-coding transcripts in a way that they will not be translated into proteins or that they will be translated into non-functional proteins. Loss-of-function variants may change one amino acid codon into a premature stop codon (nonsense variants), change the reading frame during translation (frameshift indels) or change mRNA splicing (splicing variants). As such, potentially functional variants in protein-coding regions are assumed to be easier to detect (e.g., by association analyses) than variants that moderate gene expression [5–7]. Thus, missense and LOF variants are typically prioritised as putative causal variants for the traits of interest (e.g., [8–11]).

Missense and LOF mutations can be pathogenic. For instance, missense and nonsense variants account for 57% of the entries in the Human Gene Mutation Database [12] (accessed on 30 April 2021), while small indels account for 22% and splicing variants account for another 9%. Similarly, in livestock species many missense and LOF variants have been described as causal of genetic diseases and post-natal defects ([13–16]; Online Mendelian Inheritance in Animals [17], accessed on 30 April 2021), embryonic lethality [18, 19] or product defects [20, 21]. Deleterious missense and LOF variants are subject to purifying selection and are more likely to be rare, because they are related to unfavourable phenotypes such as disease risk or reduced fertility.

However, some missense and LOF mutations can be beneficial too [22]. Moreover, some alleles that would be detrimental in the wild may be preferred in artificial selection settings. The artificial selection performed in livestock and crop breeding programs is expected to increase the frequency of alleles that favourably affect the traits included in the selection objectives. Therefore, it is also expected that missense and LOF variants are responsible for differences among breeds, genetic lines, and varieties of livestock and crop species that have undergone diverse selection histories. Identification of such functional variants would have direct applications in gene-assisted and genomic selection [23–25]. Furthermore, strategies based on genome editing have been theorized to either promote favourable alleles [26] or remove deleterious alleles [27] in selection candidates. Nevertheless, there is still limited knowledge about the existing missense and LOF variation in commercial livestock populations, in particular regarding population-specific variation, often referred to as ‘private’, and how it can affect applications such as across-breed genomic prediction.

Next-generation sequencing has great potential for livestock breeding. One of its main benefits is the power to detect large amounts of variants, many of which will be specific to the population under study. Sequencing a large number of individuals is necessary to achieve high variant discovery rates, particularly for low-frequency variants [28, 29]. There are several sequencing studies that profile the genomic variation in pigs [30–32], cattle [33, 34], or chicken [35]. These studies involved the sequencing of a low number of individuals (up to a few hundreds) at intermediate or high sequencing coverage. Alternatively, low sequencing coverage allows affordable sequencing of a much larger number of individuals, which would enable the identification of a much larger number of variants.

The objective of this study was to characterize the genetic variants detected in nine intensely selected pig lines with diverse genetic backgrounds. Particular emphasis was given to quantifying rare and population-specific functional variants, as well as the number of missense and LOF variants that an average individual carried. We also assessed the contribution of population-specific functional variants to the variance of production traits.

## Materials and Methods

### Populations and sequencing strategy

We re-sequenced the whole genome of 7,848 individuals from nine commercial pig lines (Genus PIC, Hendersonville, TN) with a total sequencing coverage of approximately 32,114x. Breeds of origin of the nine lines included Large White, Landrace, Pietrain, Hampshire, Duroc and synthetic lines. Sequencing effort in each of the nine lines was proportional to population size. The number of pigs that were available in the pedigree of each line and the number of sequenced pigs, by coverage, is summarized in Table 1. Approximately 1.5% (0.9-2.1%) of the pigs in each line were sequenced. Most pigs were sequenced at low coverage, with target coverage of 1 or 2x. A subset of pigs in each line was sequenced at higher coverage of 5, 15, or 30x. Thus, the average individual coverage was 4.1x, but the median coverage was 1.5x. The population structure across the nine lines was assessed with a principal component analysis using the sequenced pigs and is shown in Figure S1.

**Table 1.** Number of sequenced and analysed pigs.

| Line | Individuals<br>sequenced | Individuals sequenced<br>by coverage |  |  |  | Individuals used in analyses |  |  |
| --- | --- | --- | --- | --- | --- | --- | --- | --- |
|  |  | 1x | 2x | 5x | 15–30x | Pedigree | Imputed | GWAS |
| A | 1,856 | 1,044 | 649 | 73 | 90 | 122,753 | 104,661 | 88,342 |
| B | 1,491 | 628 | 728 | 54 | 81 | 84,420 | 66,608 | 56,173 |
| C | 1,366 | 685 | 545 | 44 | 92 | 88,964 | 76,230 | 64,285 |
| D | 760 | 394 | 274 | 27 | 65 | 50,797 | 41,573 | - |
| E | 731 | 362 | 311 | 16 | 42 | 79,981 | 60,474 | - |
| F | 701 | 351 | 255 | 28 | 67 | 52,470 | 39,263 | - |
| G | 445 | 217 | 176 | 15 | 37 | 21,129 | 17,224 | - |
| H | 381 | 193 | 137 | 16 | 35 | 35,309 | 29,330 | - |
| I | 321 | 111 | 158 | 18 | 34 | 15,495 | 5,247 | - |

The sequenced pigs and their coverage were selected following a three-part sequencing strategy with the objective of representing the haplotype diversity in each line. First (1), top sires and dams with the largest number of genotyped progeny were sequenced at 2x and 1x, respectively. Sires were sequenced at greater coverage because they individually contributed more progeny than dams. Then (2), the individuals with the greatest genetic footprint on the population (i.e., those that carry more of the most common haplotypes) and their immediate ancestors were sequenced at a coverage between 1x and 30x (AlphaSeqOpt part 1; [36]). The target sequencing coverage was assigned by an algorithm that maximises the expected phasing accuracy of the common haplotypes from the accumulated family information. Finally (3), pigs that carried haplotypes with low cumulated coverage (below 10x) were sequenced at 1x (AlphaSeqOpt part 2; [37]). Sets (2) and (3) were based on haplotypes inferred from marker array genotypes (GGP-Porcine HD BeadChip; GeneSeek, Lincoln, NE), which were phased using AlphaPhase [38] and imputed using AlphaImpute [39].

Most sequenced pigs, as well as pedigree relatives, were also genotyped with marker arrays either at low density (15k markers) using the GGP-Porcine LD BeadChip (GeneSeek) or at high density (80k markers) using the GGP-Porcine HD BeadChip (GeneSeek).

### Sequencing and data processing

Tissue samples were collected from ear punches or tail clippings. Genomic DNA was extracted using Qiagen DNeasy 96 Blood & Tissue kits (Qiagen Ltd., Mississauga, ON, Canada). Paired-end library preparation was conducted using the TruSeq DNA PCR-free protocol (Illumina, San Diego, CA). Libraries for resequencing at low coverage (1 to 5x) were produced with an average insert size of 350 bp and sequenced on a HiSeq 4000 instrument (Illumina). Libraries for resequencing at high coverage (15 or 30x) were produced with an average insert size of 550 bp and sequenced on a HiSeq X instrument (Illumina). All libraries were sequenced at Edinburgh Genomics (Edinburgh Genomics, University of Edinburgh, Edinburgh, UK).

DNA sequence reads were pre-processed using Trimmomatic [40] to remove adapter sequences. The reads were then aligned to the reference genome *Sscrofa11.1* (GenBank accession: GCA_000003025.6) using the BWA-MEM algorithm [41]. Duplicates were marked with Picard (http://broadinstitute.github.io/picard). Single nucleotide polymorphisms (SNPs) and short insertions and deletions (indels) were identified with GATK HaplotypeCaller (GATK 3.8.0) [42, 43] using default settings. Variant discovery was performed separately for each individual and then a joint variant set for each population was obtained by extracting the variant positions from all the individuals in it. Between 20 and 30 million variants were discovered in each population.

We extracted the read counts supporting each allele directly from the aligned reads stored in the BAM files using a pile-up function. This approach was set to avoid biases towards the reference allele introduced by GATK when applied on low-coverage whole-genome sequence data [44]. That pipeline uses pysam (version 0.13.0; https://github.com/pysam-developers/pysam), which is a wrapper around htslib and the samtools package [45]. We extracted the read counts for all biallelic variant positions, after filtering variants in potential repetitive regions with VCFtools [46]. Such variants were here defined as variants that had mean depth values 3 times greater than the average realized coverage. A total of 46,344,624 biallelic variants passed quality control criteria across all lines (see Supplementary Methods).

### Genotype imputation

Genotypes were jointly called, phased and imputed for a total of 537,257 pedigree-related individuals using the ‘hybrid peeling’ method implemented in AlphaPeel [47–49], which used all available marker array and whole-genome sequence data. Imputation was performed separately for each line using its complete multi-generational pedigree, which encompassed from 15,495 to 122,753 individuals each (Table 1). We have previously published reports on the accuracy of imputation in the same populations using this method [48]. The estimated average individual-wise dosage correlation was 0.94 (median: 0.97). Individuals with low predicted imputation accuracy were removed before further analyses. An individual was predicted to have low imputation accuracy if itself or all of its grandparents were not genotyped with a marker array or if it had a low degree of connectedness to the rest of the population. These criteria were based on the analysis of simulated and real data on imputation accuracy [48]. A total of 440,610 individuals remained, from 5,247 to 104,661 individuals for each line (Table 1). The expected average individual-wise dosage correlation of the remaining individuals was 0.97 (median: 0.98) according to our previous estimates. We accounted for the whole minor allele frequency spectrum in our analyses. However, variants with a minor allele frequency lower than 0.023 had an estimated variant-wise dosage correlations lower than 0.90 [48].

### Variant predicted consequence types

The frequency of the alternative allele was calculated based on the imputed genotypes. We defined the ‘prevalence level’ of each variant as the number of lines in which the variant segregated. To distinguish between allele frequency and prevalence level we used the terms ‘rare’ and ‘common’ to refer to variants in terms of allele frequency and ‘private’ and ‘widespread’ in terms of prevalence level, where private variants were those called only in one line and widespread variants those called in all nine studied lines. We calculated Wright’s fixation statistic (F_ST_) [50] for each variant among the lines where the variant segregated as F_ST_ = (H_T_–H_S_)/H_T_, where H_T_ is the expected heterozygosity across the combined lines assuming Hardy-Weinberg equilibrium and H_S_ is the average heterozygosity within lines assuming Hardy-Weinberg equilibrium.

Variants were annotated using Ensembl Variant Effect Predictor (Ensembl VEP; version 97, July 2019) [51] using both Ensembl and RefSeq transcript databases. For variants with multiple predicted consequence types (e.g., in the case of multiple transcripts), the most severe predicted consequence type for each variant was retrieved. Stop-gain, start-loss, stop-loss, splice donor, and splice acceptor variants were classified as LOF variants. While frameshift indels are typically included in the LOF category, we considered them as a separate category in order to quantify their impact separately. The SIFT scores [52] for missense variants were retrieved from the Ensembl transcript database. Missense variants for which SIFT scores were available were then classified either as ‘deleterious’ when their SIFT score were less than 0.05, or ‘tolerated’ otherwise. We considered the predicted consequence types of LOF, frameshift and in-frame indels, and missense as putatively functional variants. To account for the regulatory role of promoters, we classified variants within 500 bp upstream of the annotated transcription start site together with the variants in the 5’ untranslated region (UTR). This was motivated because both regions are likely to contain regulatory elements that affect transcription and because the same variant can be simultaneously in the promoter or in the 5’ UTR of different annotated transcripts for the same gene. With this action, 6.6% of the variants that were initially classified by Ensembl VEP as ‘variants upstream of gene’ were reclassified as ‘variants in promoter regions’. For further analyses, variants in promoters and in the 5’ and 3’ UTR were jointly considered (Promoter+UTR). Because some variants such as stop-gain (LOF) variants or frameshift indels are considered more likely to be benign when located towards the end of the transcripts (e.g., [53]), we analysed the relative position of these variants within transcripts (i.e., position accounting for transcript length).

### Load of putatively functional alleles

We used the imputed genotypes to estimate the number of alleles of each predicted consequence type and prevalence level that an individual carried on average. For the most common predicted consequence types, that number was estimated from 50,000 variants sampled randomly. For tolerated missense variants, we used the 50,000 variants with the highest SIFT scores. To account for the different number of variants within each predicted consequence type and prevalence level category, we calculated ‘heterozygosity’ as the percentage of variants of each category that an individual carried in heterozygosis, and the ‘homozygosity for the alternative allele’ as the percentage of variants of each category that an individual carried in homozygosis for the alternative allele.

### Association to production traits

To further explore the association of variants by prevalence level and functional annotation to selected traits, we performed genome-wide association studies (GWAS) for the three largest lines. For each line, we performed GWAS for average daily gain, backfat thickness, and loin depth using all the called variants that passed filtering (Table 2). These three traits were chosen because they are complex traits with moderate heritability estimates (range: 0.21 to 0.38). The number of pigs with records that were included in the GWAS are provided in Table 1. Most pigs with records were born during the 2008–2020 period. Breeding values were estimated by line with a linear mixed model that included polygenic and non-genetic (including contemporary group, litter, and weight as relevant for each trait) effects. Deregressed breeding values were obtained following the method of VanRaden et al. [54]. Only individuals in which the trait was directly measured were retained for the GWAS. We fitted a univariate linear mixed model that accounted for the genomic relationship as:

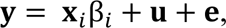

**Table 2.** Number of variants by line.

| Line | Biallelic<br>variant<br>sites (M) | SNPs |  |  | Indels |  |  |
| --- | --- | --- | --- | --- | --- | --- | --- |
|  |  | All biallelic<br>(M) | Private<br>(M) | Widespread<br>(M) | All biallelic<br>(M) | Private<br>(M) | Widespread<br>(M) |
| A | 28.83 | 24.38 | 1.56 | 8.38 | 4.44 | 0.39 | 1.56 |
| B | 28.57 | 24.32 | 2.74 | 8.38 | 4.24 | 0.51 | 1.56 |
| C | 28.88 | 24.60 | 2.51 | 8.38 | 4.28 | 0.44 | 1.56 |
| D | 21.44 | 17.94 | 1.23 | 8.38 | 3.50 | 0.32 | 1.56 |
| E | 19.06 | 15.71 | 0.51 | 8.38 | 3.35 | 0.22 | 1.56 |
| F | 20.21 | 16.86 | 0.42 | 8.38 | 3.35 | 0.16 | 1.56 |
| G | 23.38 | 19.64 | 0.50 | 8.38 | 3.74 | 0.16 | 1.56 |
| H | 22.32 | 18.78 | 0.37 | 8.38 | 3.55 | 0.12 | 1.56 |
| I | 24.59 | 20.82 | 0.76 | 8.38 | 3.77 | 0.13 | 1.56 |
| Total | 46.30 | 38.64 | 10.60 | 8.38 | 7.70 | 2.44 | 1.56 |

where 𝐲𝐲 is the vector of deregressed breeding values, 𝐱𝐱_𝑖𝑖_ is the vector of genotypes for the 𝑖𝑖th variant coded as 0 and 2 if homozygous for either allele or 1 if heterozygous, β_𝑖𝑖_ is the allele substitution effect of the 𝑖𝑖th variant on the trait, 𝐮𝐮∼𝑁𝑁(0, σ^2^ 𝐊𝐊) is the vector of polygenic effects with the covariance matrix equal to the product of the polygenic additive genetic variance σ^2^ and the genomic relationship matrix 𝐊𝐊, and 𝐞𝐞 is a vector of uncorrelated residuals. Due to computational limitations, the genomic relationship matrix 𝐊𝐊 was calculated using only imputed genotypes for the high-density marker array and its single-value decomposition was taken. We used the FastLMM software [55, 56] to fit the model.

We considered the associations with a p-value equal or smaller than 10^-6^ as significant. We calculated an enrichment score for each predicted consequence type and prevalence level category as:

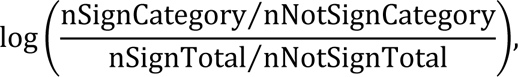

where nSignCategory was the number of variants with significant association (with at least one trait in one of the three lines) in a predicted consequence type and prevalence level category, nNotSignCategory was the number of variants with no significant association in the same category, and nSignTotal and nNotSignTotal were the total numbers of variants with and without significant association, respectively.

Linkage disequilibrium is pervasive between nearby significant variants due to the extremely high variant density of whole-genome sequence data. To account for this, we defined haplotype blocks so that only a single variant per haplotype block was considered as the putative driver of the association detected in that region. We defined the haplotype blocks for each line separately using the *--blocks* function in Plink 1.9 [57, 58]. To define haplotype blocks, pairs of variants within 5 Mb of each other were considered to be in strong linkage disequilibrium if the bottom of the 90% confidence interval of D’ was greater than 0.7 and the top of the confidence interval was at least 0.9. If the top of the confidence interval was smaller than 0.7, it was considered as strong evidence for historical recombination between the two variants. The remaining pairs of variants were considered uninformative. Regions where at least 90% of the informative pairs showed strong linkage disequilibrium were defined as a haplotype block. Within each haplotype block, we selected one ‘candidate variant’ as the variant with the most severe predicted consequence type. If there was more than one variant with the same predicted consequence type, the one with the lowest p-value was selected. This process was performed separately for each trait and line. Establishing which of the variants in linkage disequilibrium is the most likely to be causal remains one of the greatest challenges in genomics. Nevertheless, keeping the most severe variant in a haplotype block is a common assumption for prioritisation of candidate variants.

We calculated the additive genetic variance explained by each variant as 2*pq*𝛽𝛽^2^, where *p* and *q* were the allele frequencies and 𝛽𝛽 was the estimated allele substitution effect of the variant. We expressed the additive genetic variance explained by each variant as a percentage of the phenotypic variance of each trait. Finally, we calculated the median F_ST_ of the candidate variants within each predicted consequence type and prevalence level category. We compared the median F_ST_ of the candidate variants to the median F_ST_ of the same category as the logarithm of the ratio of the former to the latter.

## Results

### Prevalence of variants

A large percentage (21%) of the 46,344,624 biallelic variants that passed quality control criteria were widespread across all nine lines. Private variants represented a much smaller percentage (2 to 11%) of the variants called within each line. However, when counted across lines, private variants cumulatively predominated (28%) over the widespread ones. Most variants were neither private nor widespread. The distribution of these variants by line is shown in Table 2. Most variants (38,642,777) were SNPs, of which 10,595,681 were called in a single line (27%; 366,486 to 2,743,965 within each line) and 8,377,578 (22%) were called in all nine lines. The remaining 7,701,847 variants were indels, of which 2,436,674 were called in a single line (32%; 121,525 to 506,149 in each line) and 1,560,353 (20%) were called in all nine lines.

### Distribution of variants and relationship with recombination rate

The number of variants by chromosome was strongly correlated with chromosome length (r=0.98, P<0.05; Table S1). The average variant density by chromosome was negatively correlated with chromosome length (r=–0.87, P<0.05; Table S1). The distribution of variants within chromosomes was positively correlated to recombination rate (r=0.65, P<0.05, between variant density and recombination rate in 1-Mb non-overlapping windows [59]; Figure 1a). For example, within line A, there was on average one variant every 81 bp, but in the 5% 1-Mb windows with the lowest and highest recombination rates there was on average one variant every 152 and 54 bp, respectively (2.8-fold more variants in windows with high recombination rate). Across all lines, there was one variant every 49 bp on average, but in the 5% 1-Mb windows with the lowest and highest recombination rates there was on average one variant every 79 and 34 bp, respectively (2.3-fold more variants in windows with high recombination rate).

**Figure 1.**
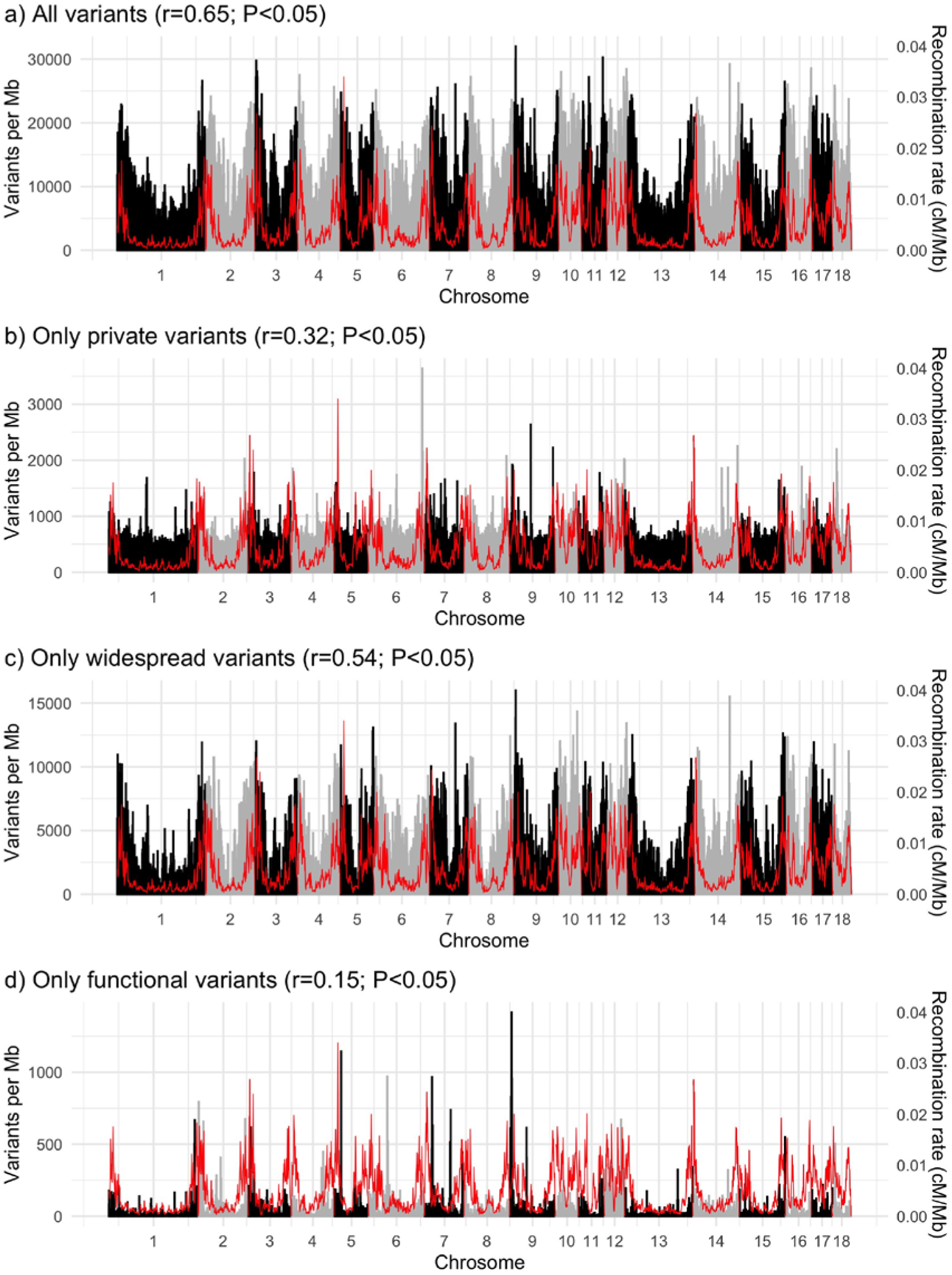
Variant density in line A (black and grey bars) and recombination rate (red line). The correlation (r) between variant density and recombination rate in 1-Mb non-overlapping windows is reported.

The distribution of private and widespread variants along the genome also differed. The distribution of widespread variants was more correlated with recombination rate than that of private variants (Figures 1b and 1c). As a consequence, private variants represented a larger proportion of the variation in regions with low recombination rate, which were depleted of widespread variants. Across all lines, in the 5% 1-Mb windows with the highest recombination rates there was on average one private variant every 167 bp and one widespread variant every 148 bp (1.1-fold more private variants relative to widespread). In the 5% 1-Mb windows with the lowest recombination rates there was on average one private variant every 260 bp and one widespread variant every 531 bp (2.0-fold more private variants relative to widespread). There were no genomic regions that were enriched for private variants across lines (Figure S2).

### Frequency and fixation index

The prevalence level and alternative allele frequency were related, in a way that less prevalent variants had also lower allele frequency (Figure 2) and lower F_ST_ (Figure 3). Private variants had an average alternative allele frequency of 0.03 (SD 0.09) as opposed to widespread variants, which had an average alternative allele frequency of 0.50 (SD 0.25). As a consequence of the less prevalent variants generally having low frequencies in the lines where they segregated, these variants showed a small degree of differentiation between the lines in which they segregated (F_ST_=0.04, SD=0.07). In contrast, the widespread variants allowed for the largest degree of differentiation between lines (F_ST_=0.21, SD=0.11).

**Figure 2.**
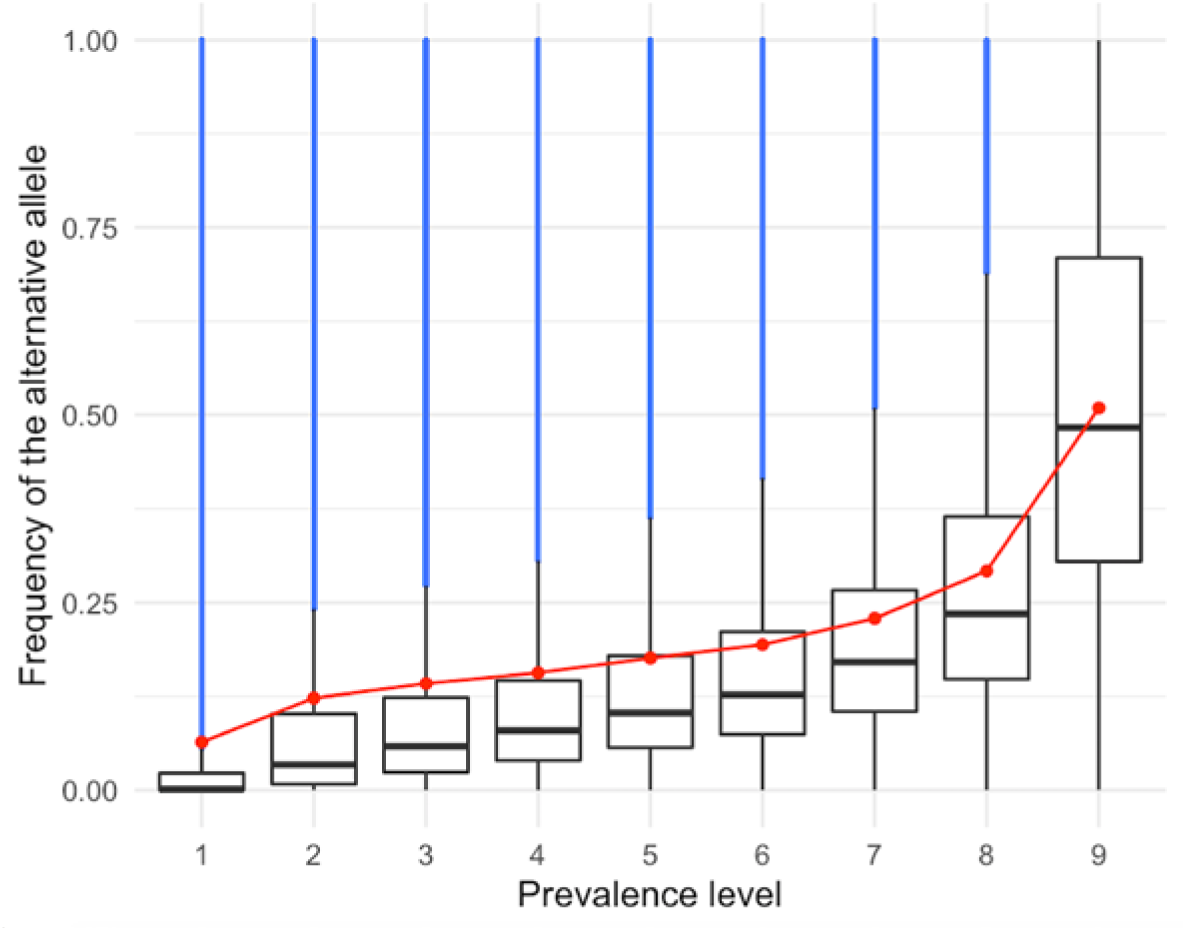
Frequency of the alternative allele by prevalence level. Red dots indicate means. In blue, values greater than 1.5 times the interquartile range.

**Figure 3.**
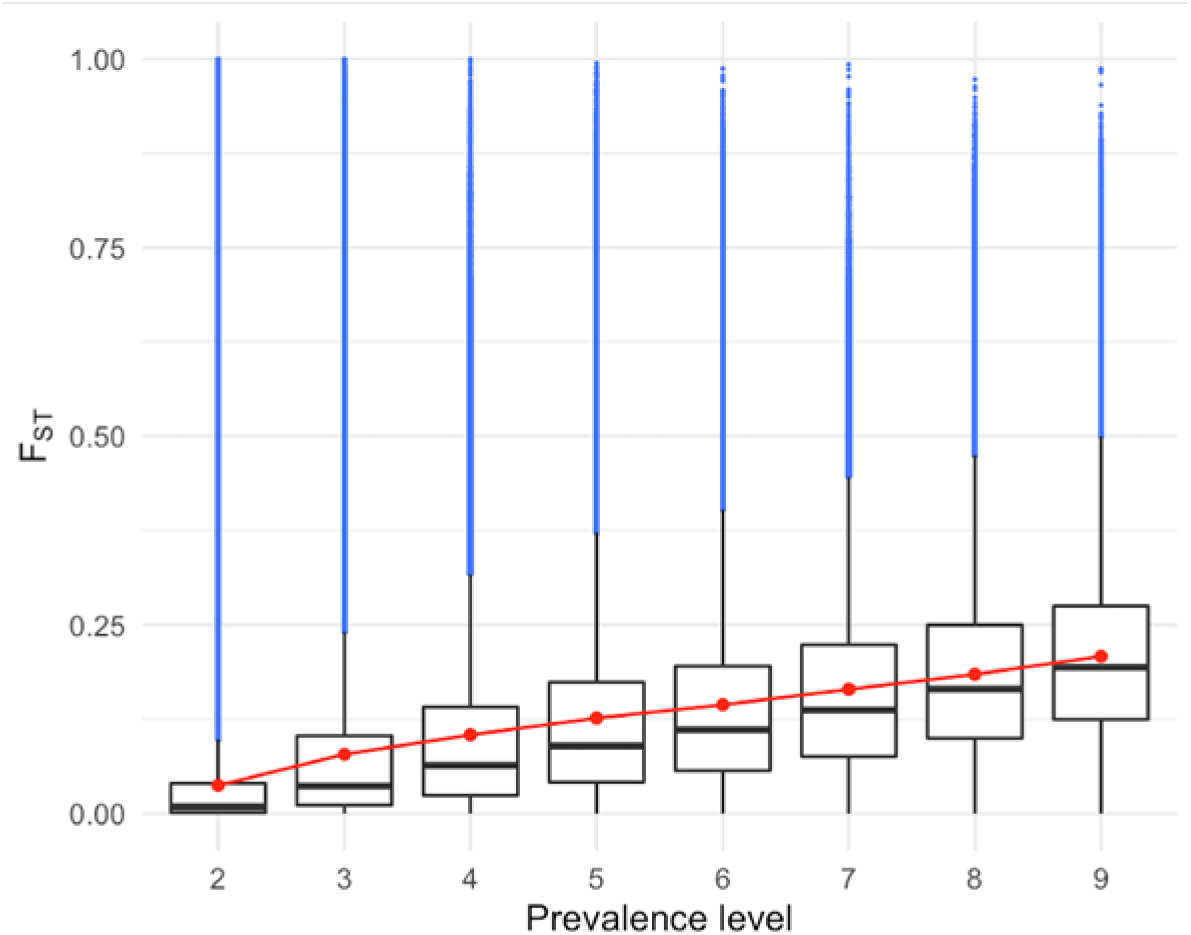
Wright’s fixation statistic (F_ST_) by prevalence level. Red dots indicate means. In blue, values greater than 1.5 times the interquartile range.

### Prevalence and frequency of putatively functional variants

The predicted consequence types of the variants are shown in Table 3. Half (49.9%) of the variants were called in intergenic regions and another 47.0% of the variants were called in intronic regions. Only 2.2% of the variants were called in the promoter or 5’ and 3’ UTR. The coding variants comprised 0.9% of the total variants, of which more than half were missense (45.5%), frameshift indels (3.1%) or LOF (3.7%). The density of putatively functional variants was only weakly correlated to recombination rate (Figures 1d).

**Table 3.** Predicted consequence types of the variants by prevalence level. The most sever consequence of each variant was used. The main Sequence Ontology (SO) terms are shown in order of severity (more severe to less severe) as estimated by Ensembl Variant Effect Predictor. The correlation (r) between the percentage of variants of each consequence type and prevalence is reported. In bold, categories that will be analysed in the next sections.

| Consequence type | Percentage of variants (%) by prevalence level |  |  |  |  |  |  |  |  |  | r |
| --- | --- | --- | --- | --- | --- | --- | --- | --- | --- | --- | --- |
|  | 1 | 2 | 3 | 4 | 5 | 6 | 7 | 8 | 9 | Total |  |
| **Loss-of-function<sup>1</sup>** | **0.061** | **0.035** | **0.026** | **0.021** | **0.019** | **0.017** | **0.019** | **0.018** | **0.019** | **0.032** | **-.76\*** |
| Splice acceptor/donor | 0.038 | 0.023 | 0.014 | 0.010 | 0.009 | 0.007 | 0.008 | 0.008 | 0.008 | 0.018 | -.79* |
| Stop-gain | 0.014 | 0.009 | 0.008 | 0.008 | 0.007 | 0.007 | 0.007 | 0.006 | 0.006 | 0.009 | -.82* |
| Stop-loss | 0.005 | 0.002 | 0.002 | 0.002 | 0.002 | 0.002 | 0.002 | 0.002 | 0.003 | 0.003 | -.36 |
| Start-loss | 0.004 | 0.002 | 0.002 | 0.002 | 0.001 | 0.001 | 0.002 | 0.002 | 0.002 | 0.002 | -.47 |
| **Frameshift indel** | **0.014** | **0.017** | **0.019** | **0.021** | **0.020** | **0.021** | **0.024** | **0.032** | **0.055** | **0.027** | **+.81\*** |
| **In-frame indel** | **0.005** | **0.008** | **0.009** | **0.008** | **0.008** | **0.008** | **0.007** | **0.007** | **0.005** | **0.006** | **-.23** |
| Missense | 0.556 | 0.378 | 0.355 | 0.340 | 0.344 | 0.336 | 0.319 | 0.306 | 0.325 | 0.393 | -.73* |
| **Deleterious** | **0.201** | **0.092** | **0.074** | **0.069** | **0.064** | **0.062** | **0.054** | **0.048** | **0.040** | **0.096** | **-.78\*** |
| **Tolerated** | **0.223** | **0.170** | **0.165** | **0.165** | **0.173** | **0.167** | **0.161** | **0.159** | **0.177** | **0.183** | **-.52** |
| Splice region | 0.105 | 0.098 | 0.088 | 0.081 | 0.083 | 0.081 | 0.080 | 0.081 | 0.085 | 0.090 | -.76* |
| **Synonymous** | **0.240** | **0.313** | **0.334** | **0.348** | **0.355** | **0.353** | **0.337** | **0.331** | **0.353** | **0.316** | **+.65** |
| **Untranslated regions** | **2.300** | **2.252** | **2.257** | **2.191** | **2.146** | **2.156** | **2.093** | **2.089** | **2.061** | **2.180** | **-.98\*** |
| Promoter + 5' UTR | 0.879 | 0.825 | 0.812 | 0.812 | 0.787 | 0.813 | 0.759 | 0.766 | 0.759 | 0.810 | -.90* |
| 3' UTR | 1.421 | 1.427 | 1.445 | 1.378 | 1.359 | 1.343 | 1.334 | 1.322 | 1.302 | 1.370 | -.94* |
| Non-coding transcript exon | 0.104 | 0.113 | 0.107 | 0.113 | 0.128 | 0.118 | 0.105 | 0.109 | 0.117 | 0.111 | +.25 |
| **Intronic** | **47.744** | **47.571** | **47.634** | **47.162** | **46.513** | **46.709** | **46.701** | **46.355** | **46.132** | **46.981** | **-.95\*** |
| Upstream of gene | 3.062 | 3.066 | 3.075 | 3.041 | 3.083 | 3.056 | 2.929 | 2.943 | 2.936 | 3.015 | -.81* |
| Downstream of gene | 2.660 | 2.679 | 2.740 | 2.747 | 2.746 | 2.705 | 2.700 | 2.707 | 2.676 | 2.692 | +.04 |
| **Intergenic** | **43.148** | **43.468** | **43.355** | **43.927** | **44.553** | **44.439** | **44.687** | **45.021** | **45.235** | **44.154** | **+.97\*** |

<sup>1</sup>If frameshift indels were included in this category: r = -.06 (P>.05)
\*Significant correlation (P<.05)

The low-prevalence variants (i.e., the variants that were identified in one or few lines) were enriched with missense and LOF variants, as well as potentially regulatory variants such as those located in the promoter and 5’ and 3’ UTR and other intronic variants. On the other hand, the high-prevalence variants (i.e., the variants that were identified in many or all lines) were enriched with frameshift indels, synonymous (non-significant correlation), and intergenic variants. Frameshift indels are typically included in the LOF category. However, our results show that the LOF category is very heterogeneous and the frameshift indels presented opposite patterns to other LOF variants. Therefore, we studied them as a separate category.

Whereas the LOF variants had lower allele frequencies than the intergenic variants in low-prevalence levels, they had similar allele frequencies in high-prevalence levels (Table 4). Thus, there was a set of LOF variants that were prevalent across lines and also had particularly high frequencies within lines. The missense variants, especially those classified as deleterious, had lower allele frequencies than the intergenic variants for all prevalence levels. The low-prevalence missense variants were enriched with a larger fraction of deleterious variants and lower SIFT scores (Figure 4). Low-prevalence stop-gain (LOF) variants and frameshift indels, unlike missense or synonymous variants, were more likely to occur towards the start of the transcripts (Figure 5). As opposed to LOF and missense variants, the frameshift and in-frame indels had intermediate allele frequencies that were much higher than those of the intergenic variants (Table 4), which indicated that in many cases the minor allele was the reference one. Within prevalence level, the LOF and deleterious missense variants had lower F_ST_ than the intergenic variants (Table 5), probably because they were kept at low allele frequencies due to negative selection pressure. The frameshift and in-frame indels also had lower F_ST_ than the intergenic variants despite their intermediate allele frequencies.

**Figure 4.**
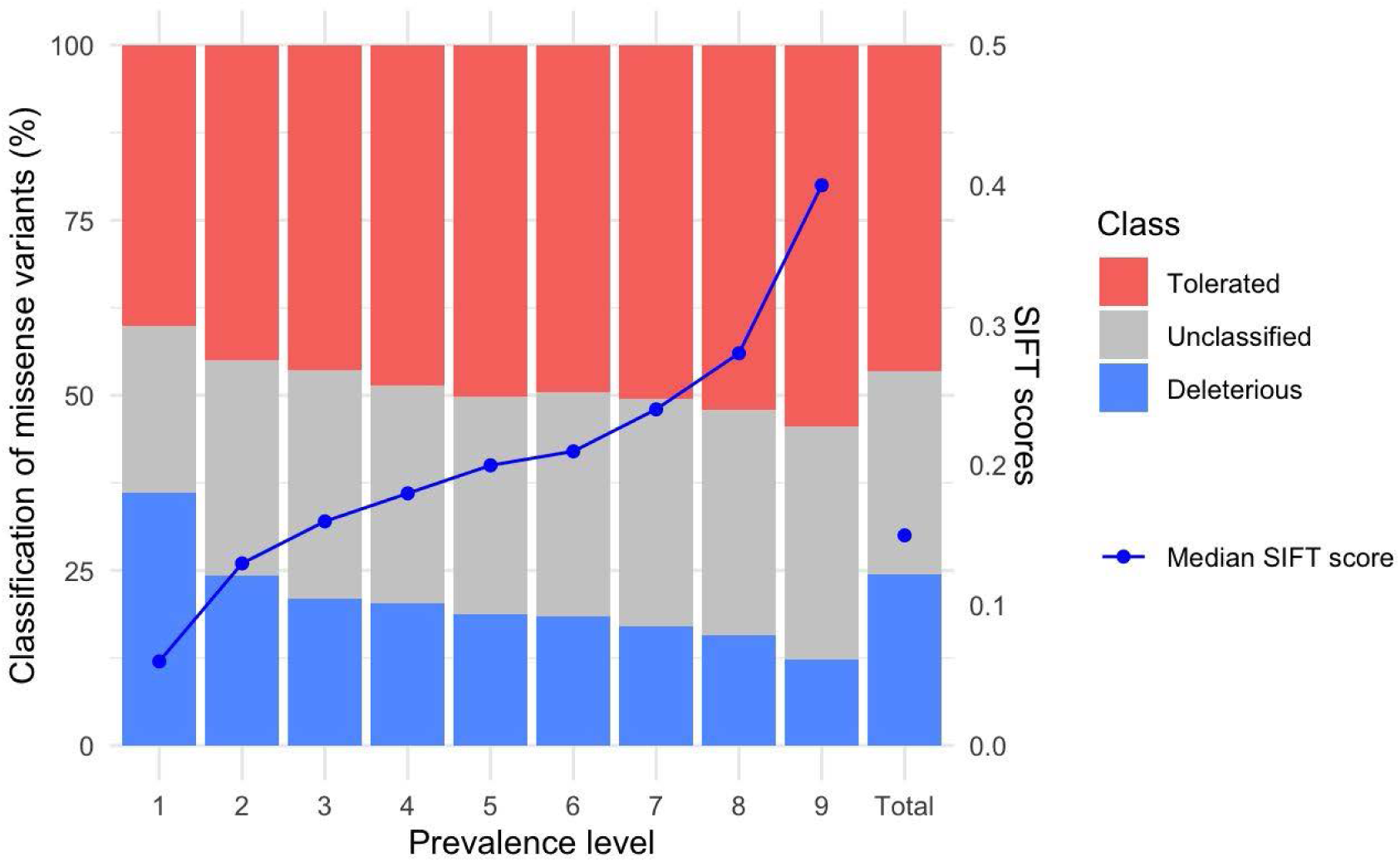
Classification of the missense variants and median SIFT score by prevalence level.

**Figure 5.**
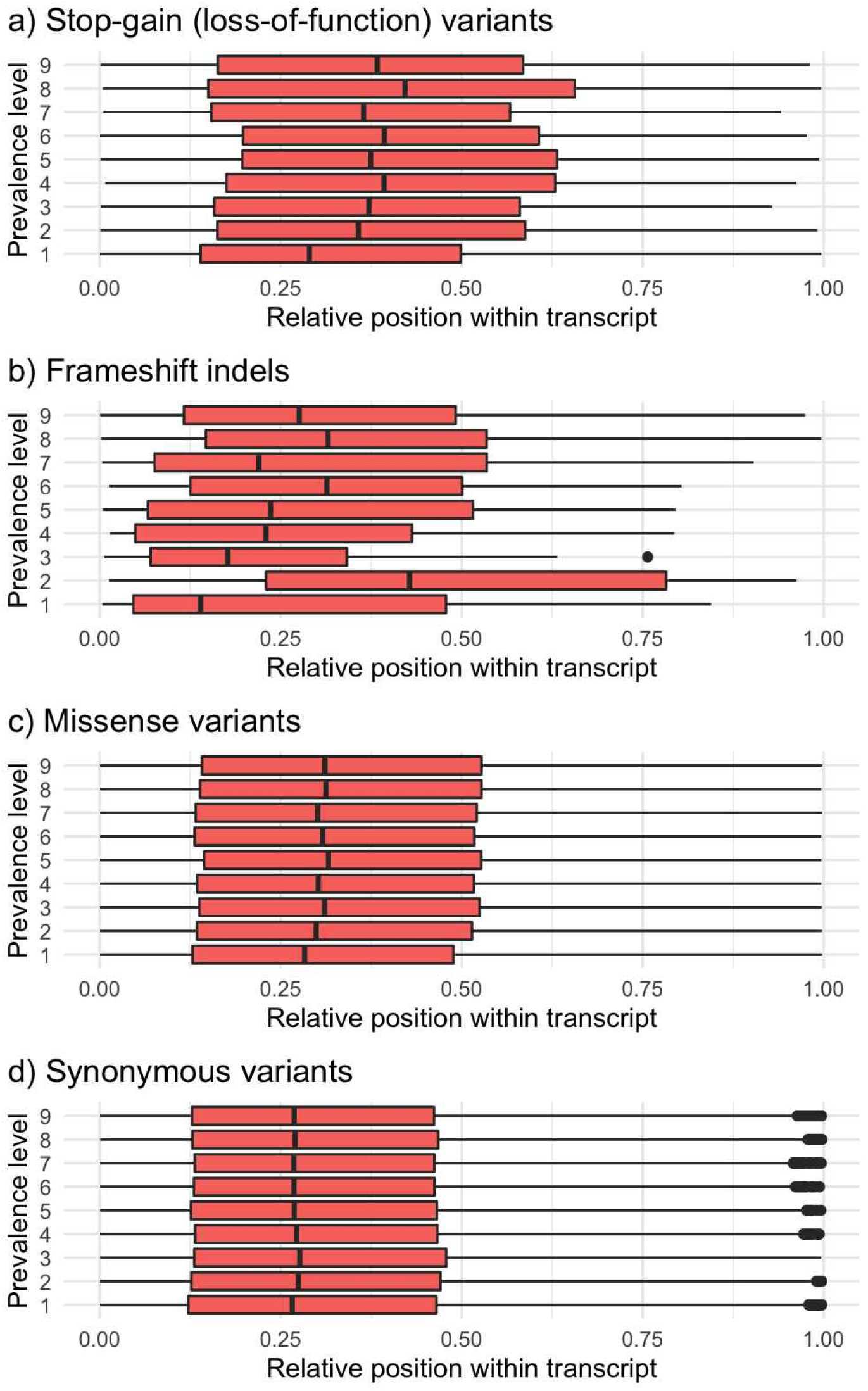
Relative position within transcript by prevalence level of stop-gain, frameshift indels, missense, and synonymous variants.

**Table 4.** Frequency of the alternative allele by predicted consequence type and prevalence level. Values are medians

| Consequence type | Frequency of the alternative allele by prevalence level |  |  |  |  |  |  |  |  |  |
| --- | --- | --- | --- | --- | --- | --- | --- | --- | --- | --- |
|  | 1 | 2 | 3 | 4 | 5 | 6 | 7 | 8 | 9 | Total |
| Loss-of-function | .0010 | .017 | .048 | .062 | .089 | .114 | .151 | .223 | .489 | .020 |
| Frameshift indel | .4816 | .758 | .757 | .420 | .302 | .260 | .339 | .456 | .693 | .634 |
| In-frame indel | .8893 | .903 | .910 | .898 | .812 | .785 | .702 | .595 | .572 | .735 |
| Deleterious missense | .0006 | .018 | .043 | .061 | .078 | .092 | .125 | .170 | .350 | .010 |
| Tolerated missense | .0011 | .027 | .047 | .066 | .083 | .106 | .143 | .202 | .443 | .074 |
| Synonymous | .0037 | .032 | .049 | .066 | .086 | .107 | .151 | .205 | .447 | .110 |
| Promoter+UTR | .0019 | .034 | .059 | .078 | .099 | .122 | .166 | .226 | .475 | .102 |
| Intronic | .0015 | .035 | .059 | .080 | .102 | .126 | .171 | .235 | .485 | .110 |
| Intergenic | .0015 | .033 | .058 | .080 | .105 | .129 | .173 | .237 | .483 | .116 |

**Table 5.** Wright’s fixation statistic (F_ST_) by predicted consequence type and prevalence level. Values are medians

| Consequence type | F<sub>ST</sub> by prevalence level |  |  |  |  |  |  |  |  |
| --- | --- | --- | --- | --- | --- | --- | --- | --- | --- |
|  | 2 | 3 | 4 | 5 | 6 | 7 | 8 | 9 | Total |
| Loss-of-function | .003 | .022 | .047 | .066 | .094 | .114 | .145 | .171 | .071 |
| Frameshift indel | .010 | .042 | .065 | .081 | .088 | .120 | .146 | .148 | .114 |
| In-frame indel | .011 | .035 | .051 | .070 | .087 | .105 | .115 | .130 | .077 |
| Deleterious missense | .005 | .029 | .055 | .073 | .087 | .110 | .131 | .160 | .068 |
| Tolerated missense | .009 | .036 | .061 | .084 | .107 | .127 | .158 | .184 | .108 |
| Synonymous | .013 | .040 | .062 | .090 | .110 | .130 | .158 | .194 | .117 |
| Promoter+UTR | .009 | .036 | .060 | .086 | .108 | .131 | .158 | .190 | .110 |
| Intronic | .009 | .037 | .063 | .089 | .111 | .136 | .164 | .195 | .118 |
| Intergenic | .009 | .036 | .066 | .091 | .112 | .139 | .167 | .193 | .121 |

### Load of putatively functional alleles by prevalence level

Most of the missense deleterious and LOF variants that an individual carried in homozygosis for the alternative allele were high-prevalence variants. Only a small proportion of these variants were private. An individual carried on average 1,048 (SD 57) LOF variants in homozygosis for the alternative allele, of which 713 (SD 36) were widespread across all nine lines and only 20 (SD 7) were private. An average individual carried 1,379 (SD 165) deleterious missense variants in homozygosis for the alternative allele, of which 1,012 (SD 79) were widespread and only 4 (SD 3) were private. An average individual carried 1,080 (SD 89) LOF and 2,632 (SD 235) deleterious missense variants in heterozygosis.

We found signals of negative selection against deleterious missense variants, in particular the private ones. Individuals proportionally carried less deleterious missense variants in homozygosis for the alternative allele than variants of other predicted consequence types, regardless of prevalence level (Figure 6). Individuals also carried proportionally less private tolerated missense, synonymous and LOF variants in homozygosis for the alternative allele than expected, but not in heterozygosis.

**Figure 6.**
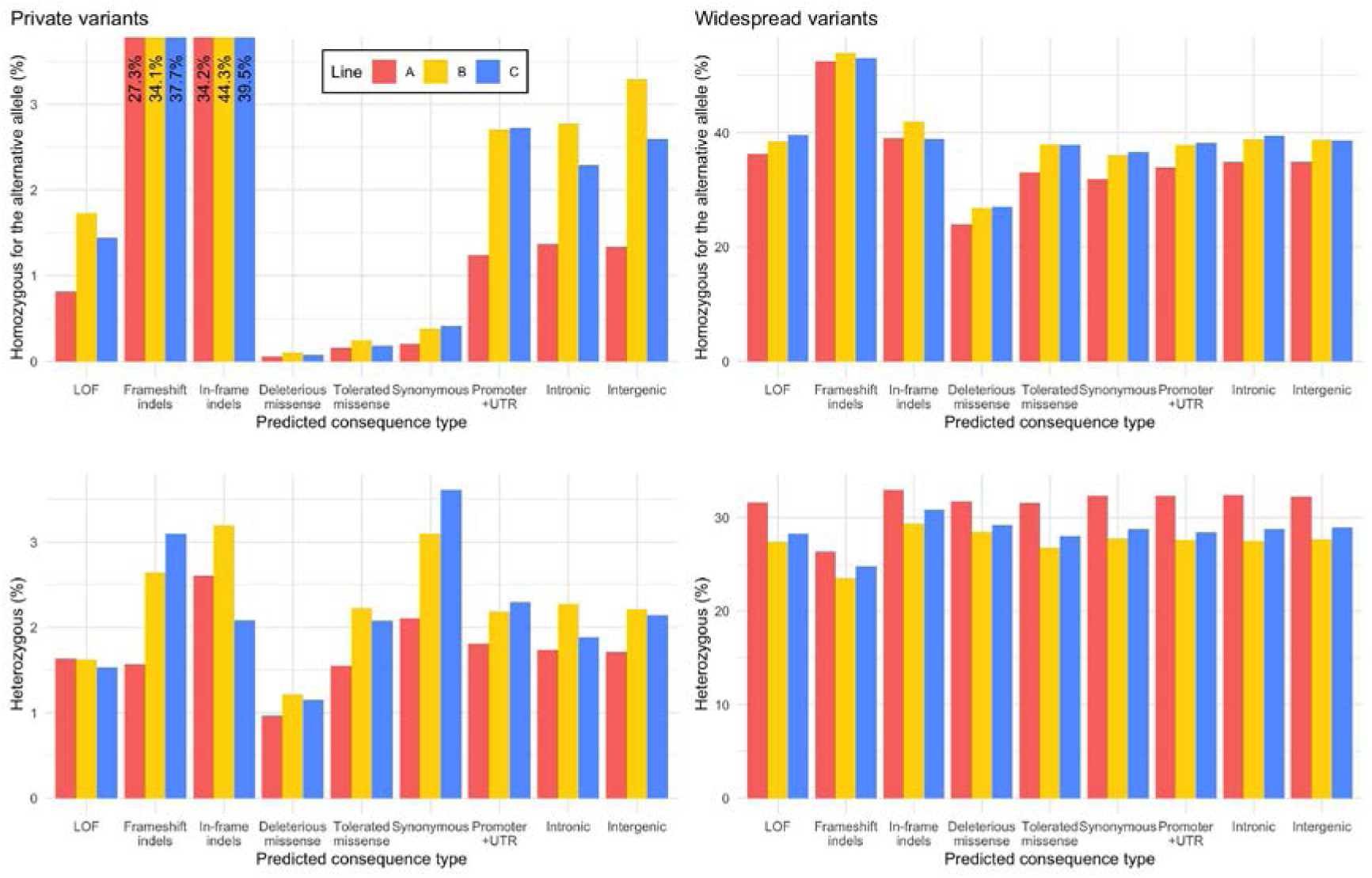
Percentage of variants in homozygosis for the alternative allele or in heterozygosis in an average individual by predicted consequence type and prevalence level. LOF: loss-of-function; UTR: untranslated regions.

### Association of low-prevalence variants to production traits

Significant variants were enriched with putatively functional and regulatory variants of different prevalence levels, and depleted of intergenic variants. A total of 108,109 variants were significantly associated to at least one trait in one line. Figures 7a and 7b summarise the enrichment scores for all significant variants. The predicted consequence types that reached the greatest enrichment scores were LOF, frameshift indels, and unclassified missense variants, with various prevalence levels. Variants with intermediate prevalence levels were amongst the most enriched. These trends were accentuated after selecting candidate variants from haplotype blocks. In each line we defined from 1,554 to 2,118 haplotype blocks. A total of 6,692 candidate variants remained after accounting for linkage disequilibrium within each haplotype block for all lines and traits. Figures 7c and 7d summarise the enrichment scores for the candidate variants. The enrichment scores based on the candidate variants revealed a stronger depletion of intergenic variants, as well as intronic (with the exception of high-prevalence), and a much stronger enrichment for LOF, frameshift indels, and missense variants. For putatively functional variants, there were no clear trends of their enrichment scores across prevalence levels. The trends of the enrichment scores between predicted consequence types and prevalence levels were similar in the three tested traits.

**Figure 7.**
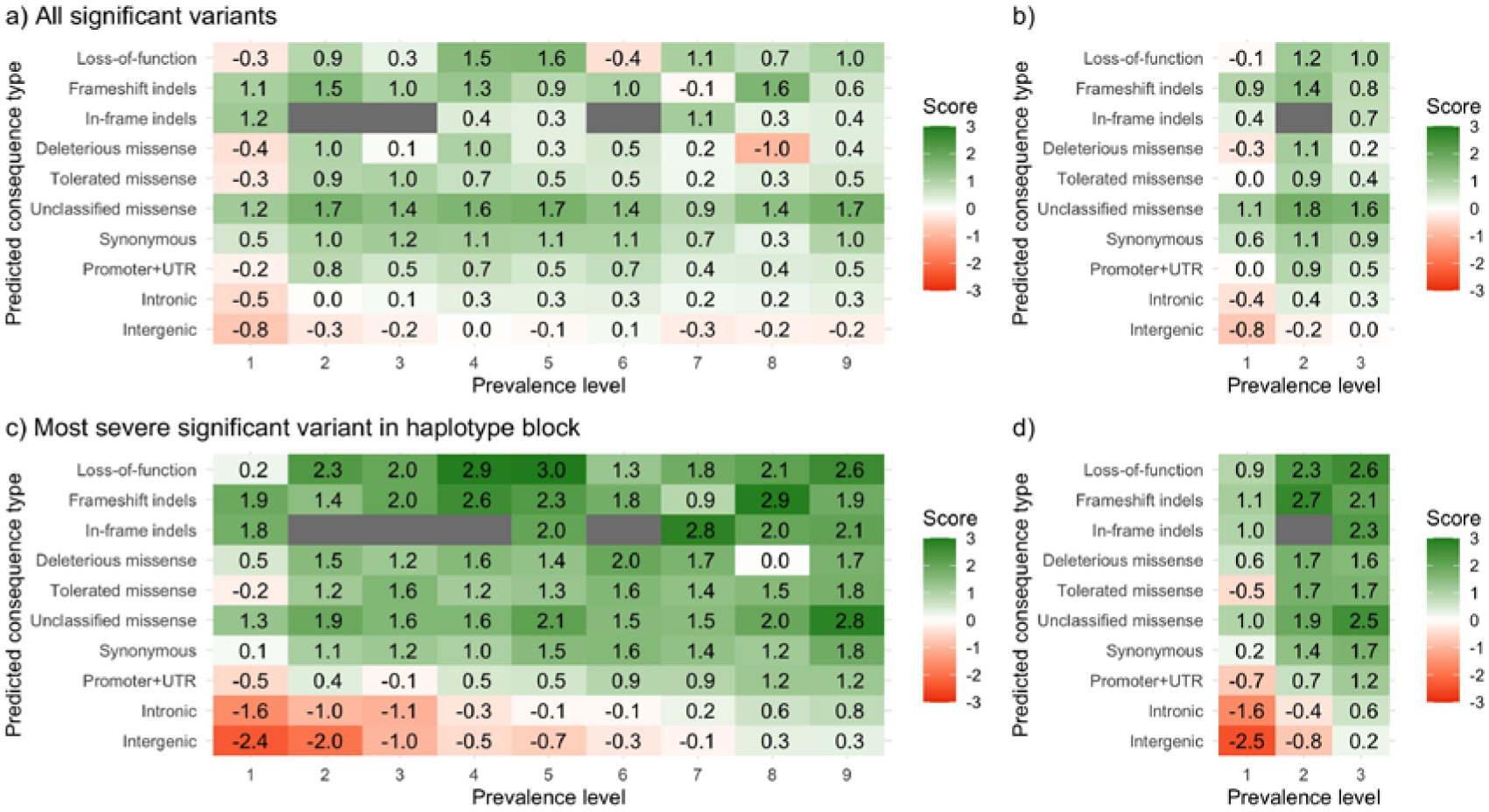
Enrichment scores for the significant variants in the genome-wide association study by variant prevalence level and predicted consequence type. Either all significant variants (panels a and b) or only the most sever significant variants within haplotype block (panels c and d) were used. Prevalence level was considered across all 9 lines (panels a and c) or only across the 3 lines included in the genome-wide association study (panels b and d).

In general, the lower allele frequency of low-prevalence variants hindered the detection of significant associations for these markers. Low-prevalence variants that were detected as significantly associated to the production traits actually had intermediate allele frequencies that were greater than expected for their prevalence level. Low-prevalence variants in general explained low percentages of variance (Figure 8), although there were some instances of low-prevalence variants that explained up to 3.2% of phenotypic variance. Significant variants had higher F_ST_ than other variants of the same predicted consequence type and prevalence level (Figure 9). This enrichment was especially strong for low-prevalence variants, which in some instances reached F_ST_ around 0.15.

**Figure 8.**
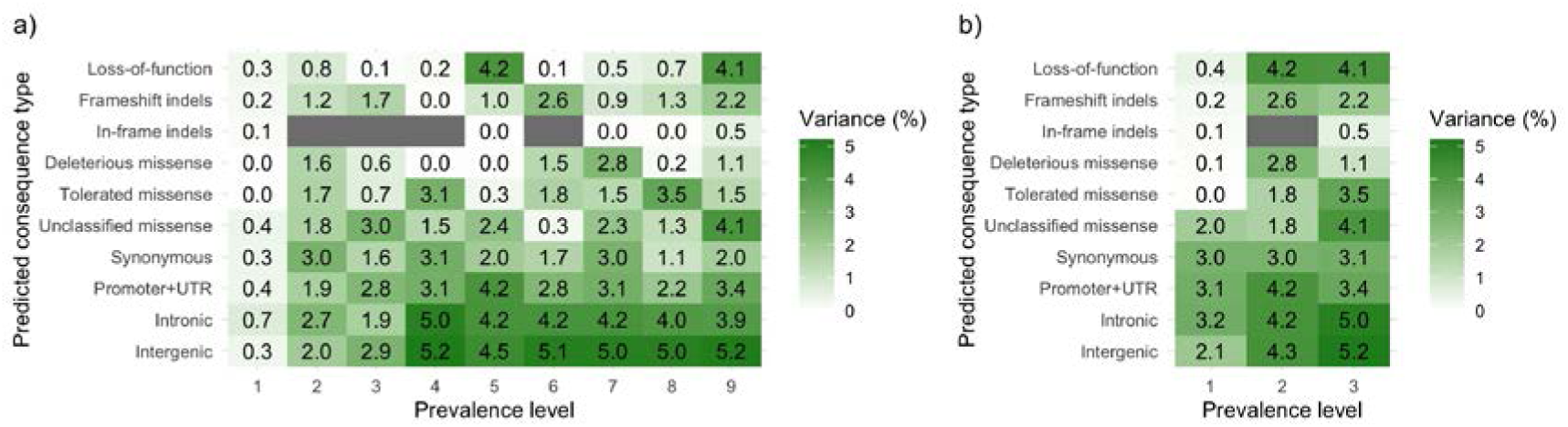
Maximum percentage of phenotypic variance explained by the individual candidate variants within each prevalence level and predicted consequence type. Only the candidate variants after accounting for linkage disequilibrium were used. Prevalence level was considered across all 9 lines (panel a) or only across the 3 lines included in the genome-wide association study (panel b).

**Figure 9.**
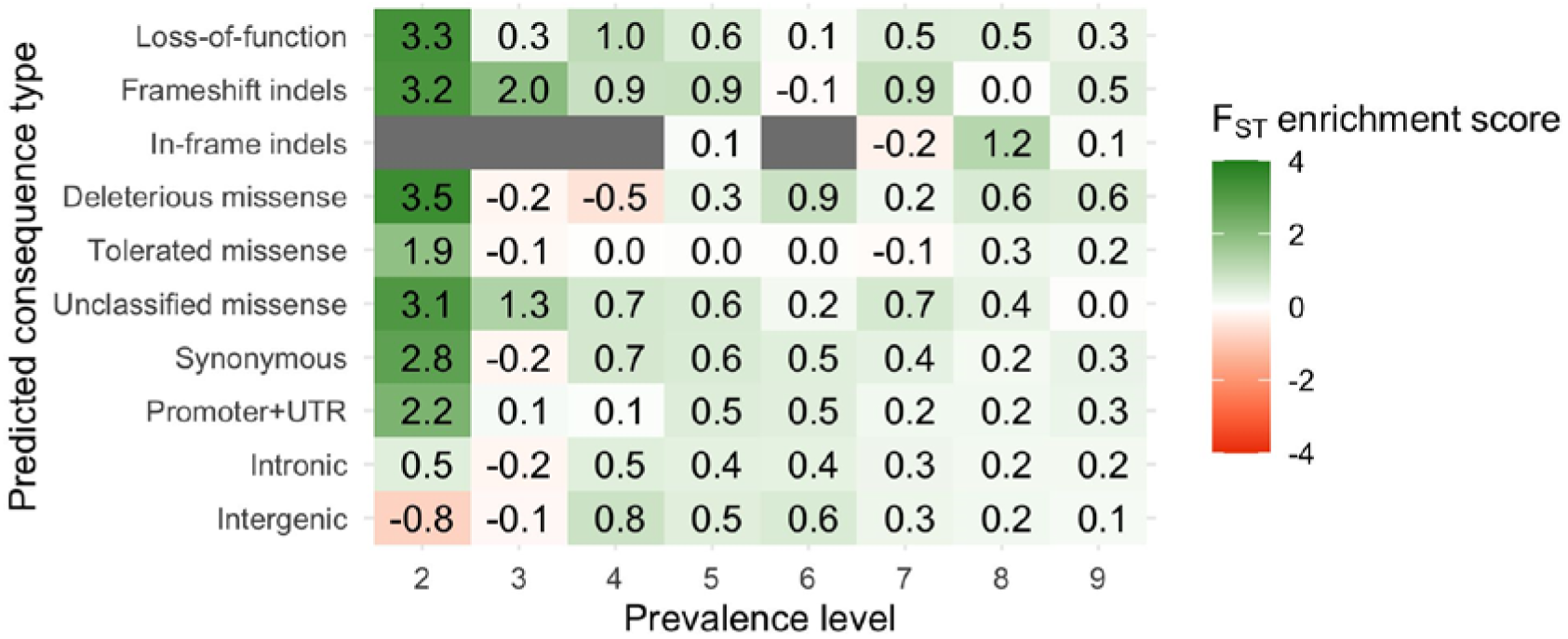
Enrichment scores for the F_ST_ median of the candidate variants within each prevalence level and predicted consequence type. Only the candidate variants after accounting for linkage disequilibrium were used. Prevalence level was considered across all 9 lines.

## Discussion

Our results contextualize the importance of population-specific and low-prevalence genetic variants. Next, we will discuss: (1) the distribution and functional annotation of low-prevalence variants, (2) the load of putatively functional alleles by prevalence level, and (3) the association of low-prevalence variants to production traits.

### Distribution and functional annotation of low-prevalence variants

The main difficulty for the study of low-prevalence genetic variants is that the prevalence of a variant across several lines is strongly related to its allele frequency, in a way that the low-prevalence variants are also rare within the lines where they occur. This is possibly because low-prevalence variants are relatively recent or are constrained by negative selection.

On one hand, the distribution of private variants was only weakly correlated to recombination rate and, therefore, regions with low recombination rate were enriched for private variants. Although the interplay between recurring sweeps, background selection and other phenomena at play is not fully understood yet, it is generally accepted that selection on linked variants leads to loss of variation in regions with low recombination rates [60]. Our observation that regions with low recombination rate were enriched for private variants suggests that private variants may have been less affected by selective sweeps than widespread variants. This would be consistent with previous observations of the younger age of rare and low-prevalence variants [61], and suggests that many private variants arose more recently in time than widespread variants, likely after line differentiation, and accumulated in low-recombining regions due to the reduced efficacy of purifying selection in those regions [62, 63].

On the other hand, the low-prevalence variants were enriched for putatively functional variants and with signs of a greater severity (stop-gain and frameshift indels that occur earlier in the transcript, and missense variants predicted to be deleterious). Variants that affect performance traits or that cause a detrimental condition are under the action of directional selection and are therefore driven towards loss or fixation [64, 65]. The low F_ST_ estimates for the low-prevalence variants indicated that selection pressure keeps these variants at low minor allele frequency even when they occur in several lines, especially if they are putatively functional [66]. This could be caused by natural selection or similar selection objectives across livestock populations. These observations were also consistent with previous reports showing that some putatively functional variant categories (such as stop-gain and deleterious missense) were enriched for variants that were private to single cattle breeds [33], that putatively functional variants were less likely to have high frequency of the alternative allele across multiple chicken lines [35], and that population-specific variants in non-African humans were enriched with putatively functional variants [67].

The relationship between variant prevalence across lines and allele frequency highlighted the suitability of using a low-coverage sequencing approach to study this fraction of genetic variation. Nonetheless, bioinformatics pipelines for calling, genotyping and, even imputing such variants should account for the increased uncertainty associated to their low allele frequency. We decided on using a very relaxed variant calling strategy with little filtering to account for as many rare variants as possible, but a sizeable fraction of these rare variants were discarded after imputation because they were fixed for the imputed individuals that passed quality control. Low-coverage sequencing is also unsuitable for other types of genetic variants, such as structural variations (CNVs, tandem duplications, and inversions), which could also be putatively functional and population-specific [68]. Of course, the number of called variants and the proportion that were private or widespread depends of the number of sequenced lines [32, 35] as well as the sequencing effort in each line. Our results also suggest that what is typically grouped as LOF is actually a heterogeneous category. In particular, frameshift indels showed patterns that did not conform to the other predicted consequence types.

### Load of putatively functional alleles by prevalence level

We found that an average individual carried a larger number of LOF and missense deleterious variants than previously reported in other livestock species or in humans. However, there is not yet a clear consensus on the number of LOF and deleterious missense alleles that are present in the genome of an average individual. In humans, it has been estimated that an average individual carries 100-150 LOF alleles [64,69–71] and around 800 weakly deleterious mutations [72], most of which are rare. In domestic livestock populations, the number of LOF and deleterious alleles carried on average by individuals has been reported to be greater than in wild populations [73], including estimates of 100 to 300 deleterious variants in domestic pigs [74], over 400 deleterious variants in domestic chicken [74], and 1,200-1,500 deleterious variants in domestic yak [75]. Similar magnitudes have been reported in dogs [76], rice [77], and sunflower [63].

It has been debated why healthy individuals carry a larger number of LOF variants in homozygosis than expected [78, 79]. These could be driven by the fact that not all predicted LOF variants are detrimental and their functional impact should be validated before being considered as such. Many predicted LOF variants are in fact neutral, advantageous (either in the wild or in controlled production environments), or even may arise simply because of sequencing and annotation errors [78]. This claim is supported by the large proportion of LOF observed in homozygosis for the alternative allele compared to the other consequence types, which casts doubt on the real impact of those variants. On the contrary, individuals carried a lower proportion of alleles predicted to be deleterious missense in homozygosis, which supports that variants predicted as such may have a real impact on genetic variation of production traits and, therefore, be subjected to selection pressure.

These observations have implications for the identification of variants to be used for genomic prediction or genomic edition strategies such as PAGE [26] or RAGE [27]. Efforts to promote or remove alleles should target variants that make a substantial contribution to traits of interest, namely functional variants. However, it is hard to computationally predict and statistically estimate the effects of such variants, especially if they have low allele frequency. The number of LOF variants in homozygosis for the alternative allele suggests that predicted loss of function is not a good indicator that a variant is strongly deleterious in the context of livestock breeding. Similarly, bioinformatics predictors of missense variant effects appear to be not very accurate [80, 81]. High-throughput fine-mapping and variant screening would be needed to ascertain variant causality and disentangle causality from linkage disequilibrium.

### Association of low-prevalence variants to production traits

Genome-wide association studies on three polygenic traits of economical importance in the three largest lines revealed that the significant markers were enriched for putatively functional roles, such as LOF, frameshift indels and missense variants, and depleted of intergenic variants. This pattern of enrichment was similar to previous reports from human datasets [82]. However, only a few of the population-specific and low-prevalence variants were significantly associated to the traits, even after accounting for linkage disequilibrium. Most of the significant variants showed intermediate or high prevalence levels. These observations are consistent with previous meta-analyses in cattle that showed that significant variants are often common variants [83]. This could be explained by either the fact that quantitative trait nucleotides have intermediate or high allele frequencies or the fact that most studies are underpowered to map rare causal variants. The latter scenario still seems more likely given that the significant private and low-prevalence variants had intermediate allele frequencies. This also supports that these significant variants have biological functions that contributed to trait phenotypic variance rather that they reached intermediate allele frequencies by drift or by hitchhiking with linked variants under selection [84]. However, these amounted to a small number of variants that explained small fractions of variance. Other more widespread variants, including intergenic variants, successfully acted as tag variants for them and captured much larger fractions of trait variance. This makes them more suitable for applications in animal breeding, as is already the case with marker arrays. A similar result was found in cattle, where splice site and synonymous variants explained the largest proportions of trait variance, while missense variants explained almost null variance [85]. It is worth pointing out that even a variant with a large allele substitution effect will explain a small percentage of variance if the alternative allele is rare.

It can be hypothesized that some of the low-prevalence variants with low allele frequency have non-negligible effects for traits of interest. Despite the large amount of individuals included in this study, the large volume of variants and the pervasiveness of linkage disequilibrium among them still make it very challenging to disentangle their contribution to trait variance. While genome-wide association studies involving more than one breed typically find multiple breed-specific associations (e.g., [86]), based on our results it seems unlikely that breed-specific associations arise from the low-prevalence variants. They would instead stem from differences in allele frequency, linkage disequilibrium structure or genetic background that affect the power to detect the effect of prevalent variants across different populations. Significant variants were enriched with higher F_ST_ estimates than non-significant variants, which is also consistent with previous reports [83]. Although the enrichment was greater for low-prevalence variants, it remains unclear to which degree these variants could relate to selection history or explain differences among lines for the studied traits.

## Conclusion

Low-prevalence variants are enriched for putatively functional variants, including LOF and deleterious missense variants. However, most low-prevalence variants are kept at very low allele frequency. Only a small subset of low-prevalence variants was found at intermediate allele frequencies and had large estimated effects on production traits. Population-specific variants that were significantly associated to complex traits had greater degrees of differentiation than non-significant variants in the same category. However, more widespread variants, including intergenic variants, successfully captured larger fractions of trait variance. Therefore, overall, not accounting for population-specific and other low-prevalence variants is unlikely to hinder across-breed analyses, such as the prediction of genomic breeding values using reference populations of a different genetic background.

## Ethics approval and consent to participate

The samples used in this study were derived from the routine breeding activities of PIC.

## Consent for publication

Not applicable.

## Availability of data and material

The software packages AlphaPhase, AlphaImpute, and AlphaPeel are available from https://github.com/AlphaGenes. The software package AlphaSeqOpt is available from the AlphaGenes website (http://www.alphagenes.roslin.ed.ac.uk). The datasets generated and analysed in this study are derived from the PIC breeding programme and not publicly available.

## Authors’ contributions

RRF, MJ, and JMH designed the study; RRF and MJ performed the analyses; RRF and MJ wrote the first draft; BDV, CYC, WHO, GG, and JMH contributed to the interpretation of the results and provided comments on the manuscript. All authors read and approved the final manuscript.

## Acknowledgements

This work has made use of the resources provided by the Edinburgh Compute and Data Facility (ECDF) (http://www.ecdf.ed.ac.uk/).

## Additional files

### Supplementary Methods

A total of 70,739,387 variants were called across all nine lines. Of these, 24,394,763 variants failed to meet quality control criteria. Of these, 148,825 variants were discarded because they had mean depth values 3 times greater than the average realized coverage, 1,927,221 were multiallelic within line, and 1,673,219 were biallelic within line but multiallelic when all lines were considered. The remaining variants were imputed for all pedigreed individuals, but 20,645,588 of them were fixed for the reference allele in the imputed individuals that passed our accuracy quality control. This affected mostly variants that had been called in only one line and for which the alternative allele segregated at very low frequency. The hypothesis that such variants arise from false positives in variant calling seems unlikely to be the main cause as for more than 99% of these variants we read the alternative allele in at least two individuals. Additionally, we previously quantified that 96.9% of the variants called from low-coverage data were confirmed by sequencing the same individuals at high coverage [1]. A total of 46,344,624 biallelic variants passed quality control criteria across all lines.

---

**Figure S1.**
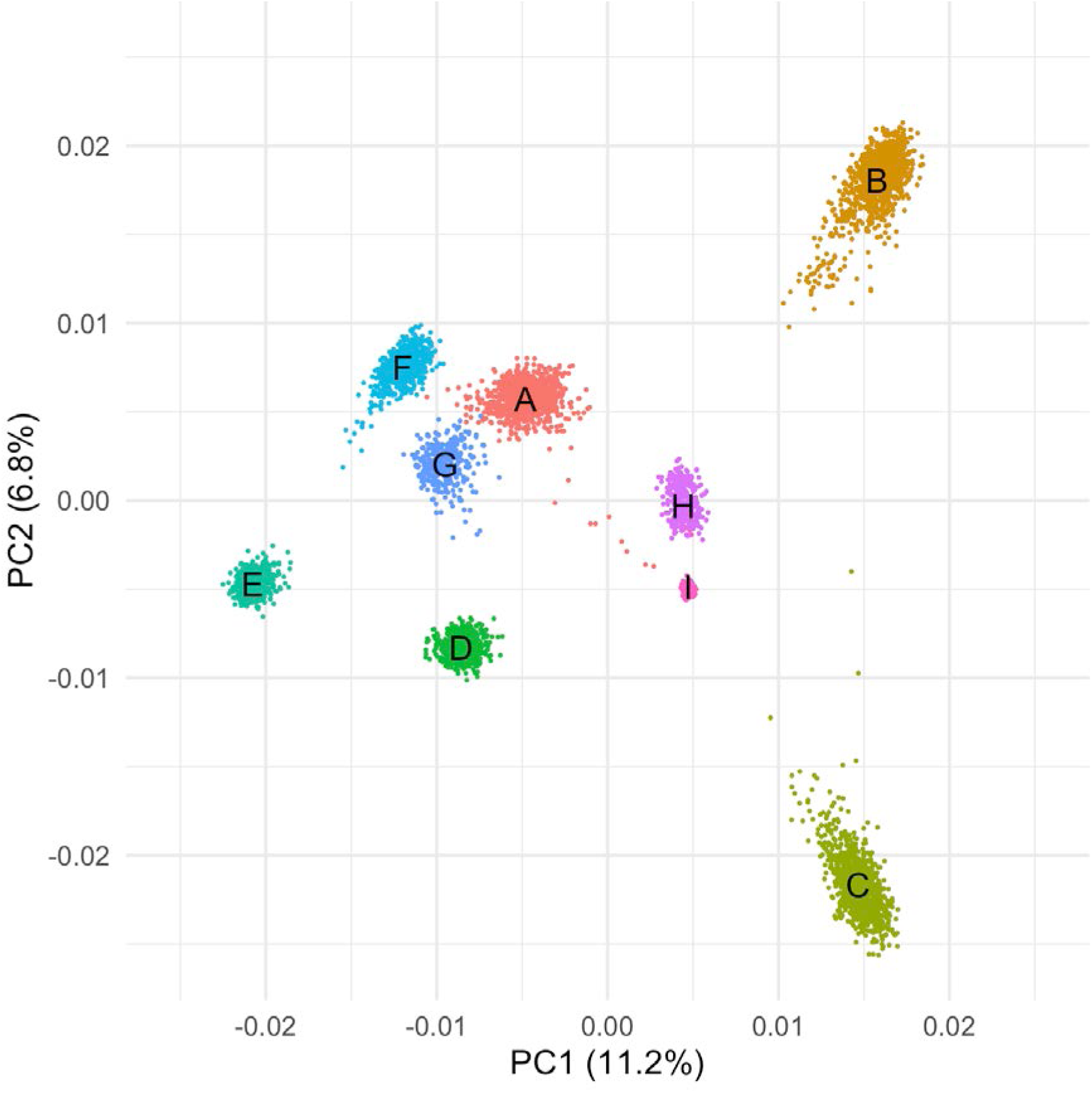
Population structure of the sequenced pigs according to the two first principal components. The colour clusters correspond to lines A to I.

**Figure S2.**
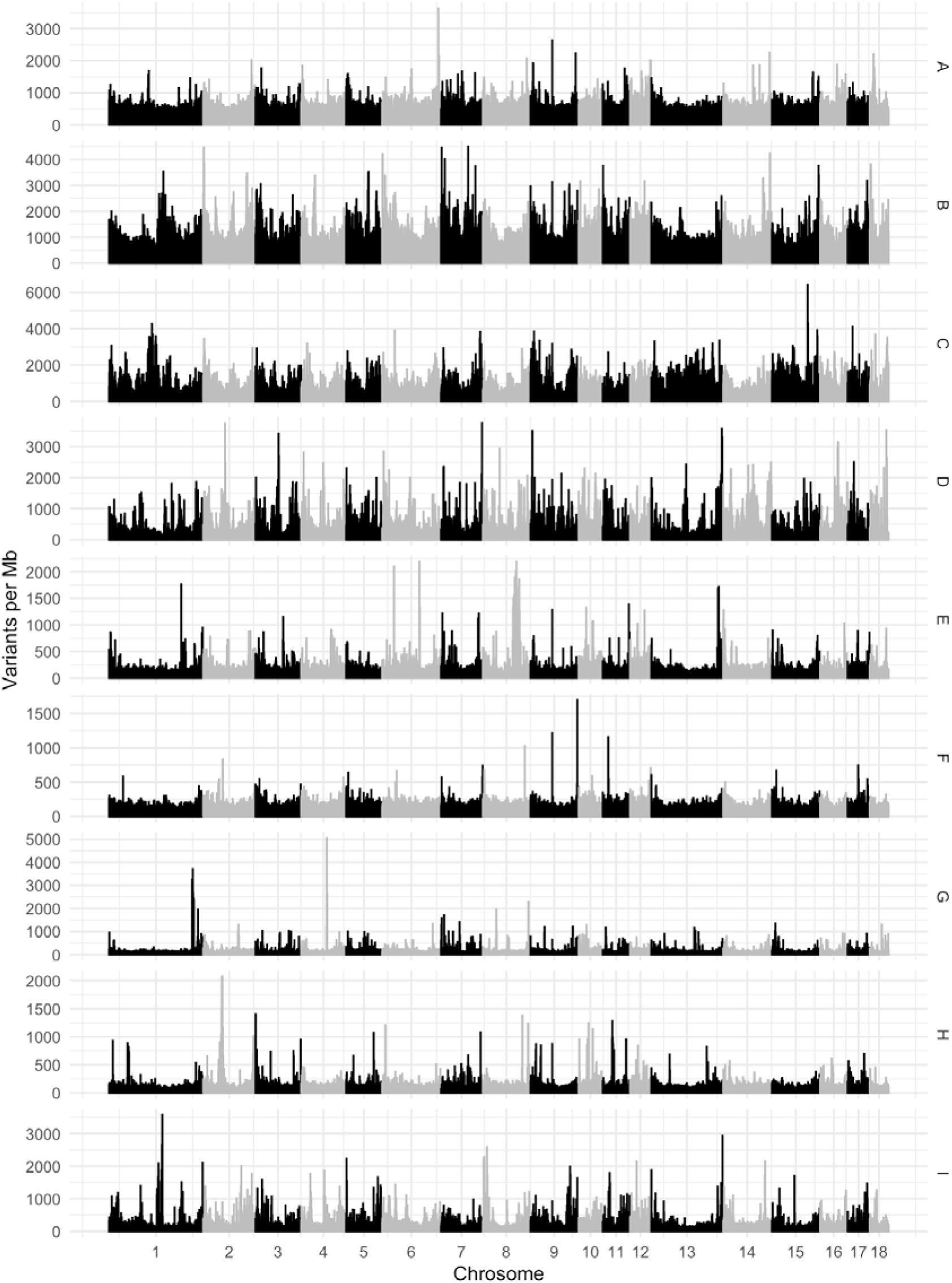
Variant density for the private variants in each line.

**Table S1.** Number of analysed variants by chromosome

| Chromosome | Length (Mb) | SNPs (M) | Indels (M) | Variant density<br>(thousands/Mb) |
| --- | --- | --- | --- | --- |
| 1 | 274.3 | 3.77 | 0.76 | 16.5 |
| 2 | 151.9 | 2.60 | 0.52 | 20.5 |
| 3 | 132.8 | 2.35 | 0.44 | 21.0 |
| 4 | 130.9 | 2.21 | 0.43 | 20.2 |
| 5 | 104.5 | 1.95 | 0.39 | 22.4 |
| 6 | 170.8 | 2.80 | 0.55 | 19.6 |
| 7 | 121.8 | 2.20 | 0.43 | 21.6 |
| 8 | 139.0 | 2.37 | 0.50 | 20.6 |
| 9 | 139.5 | 2.47 | 0.48 | 21.1 |
| 10 | 69.4 | 1.60 | 0.31 | 27.5 |
| 11 | 79.2 | 1.57 | 0.31 | 23.7 |
| 12 | 61.6 | 1.35 | 0.25 | 26.0 |
| 13 | 208.3 | 2.97 | 0.64 | 17.3 |
| 14 | 141.8 | 2.38 | 0.48 | 20.2 |
| 15 | 140.4 | 2.20 | 0.46 | 18.9 |
| 16 | 79.9 | 1.50 | 0.30 | 22.5 |
| 17 | 63.5 | 1.32 | 0.25 | 24.7 |
| 18 | 56.0 | 1.04 | 0.19 | 22.0 |
| Total | 2,501.9 | 38.64 | 7.70 | 18.5 |

